# To slide or not to slide, that is the question: evaluating dense semilandmarks and sliding in 3D geometric morphometrics with real and simulated data

**DOI:** 10.64898/2026.08.28.747867

**Authors:** A. Murat Maga

## Abstract

Dense semilandmarks describe 3D surfaces with hundreds to thousands of points, and sliding them by bending energy or Procrustes distance is a near-universal default. Three questions remain open: does dense sampling add shape beyond fixed landmarks, how many points are needed, and does sliding help or harm? Real specimens cannot answer them: the true correspondence is unknown. We tested two workflows, ALPACA (single-template registration) and DeCAL (landmark-anchored correspondence), on 496 mouse skulls at 250–1,000 points, with and without sliding, scored by surface reconstruction. We repeated it on 500 synthetic skulls with exact correspondence, measuring each point’s distance to its true homologue.

Dense semilandmarks lowered error for almost every specimen; the fixed landmarks added little but supplied anchoring the semilandmarks could not, and the anchored method was more accurate. The benefit saturated near 250 points for ALPACA but kept improving to 1,000 for DeCAL. Procrustes- distance sliding harmed every configuration; bending-energy sliding helped only a poor, landmark-free correspondence, vanishing once anatomical anchors spanned the form. Match the sliding decision to the correspondence in hand: relax a poor one, leave a good one alone, never slide toward the mean. Known-correspondence specimens offer a general test of landmarking and sliding against ground truth.

## 1. Introduction

Geometric morphometrics describes biological shape from the coordinates of homologous landmarks, points placed at anatomically corresponding locations on every specimen [1]. This works well where such points exist, but many structures have too few of them. The cranial vault, the articular surfaces, and the broad areas of bone between sutures are mostly smooth, with few discrete, repeatable features to landmark. Semilandmarks extend the method to these curves and surfaces. They are points spread along an outline or across a surface patch that stand in for the form between the anatomical landmarks [2,3]. A semilandmark’s position along the curve or surface is not anatomically defined. You could place it a little to either side without any loss, so its position is arbitrary and not homologous from one specimen to the next. In the standard modern account, only the coordinate orthogonal to the surface is anatomically meaningful; the tangential position is estimated, not measured [4]. Sliding removes this arbitrariness. It lets each semilandmark move tangentially, along the curve or surface, until the whole configuration is optimal by some criterion. The classic criterion is the bending energy of the thin-plate spline that maps each specimen to a reference [2,3]; an alternative is the Procrustes distance among specimens [5]. The aim is to turn arbitrarily spaced points into corresponding ones, so that a given semilandmark marks “the same place” on every specimen.

Three-dimensional surfaces are much harder to landmark than two-dimensional outlines. A surface patch needs hundreds to thousands of semilandmarks, and placing them by hand on the tens or hundreds of specimens a modern study needs is not feasible. So the field uses automated methods that sample the anatomy densely and quickly, by transferring a dense point set from a template onto every specimen [6–10]. Practical guides recommend sampling a whole cranium with several hundred to a thousand surface semilandmarks and, by default, sliding them to minimise bending energy [11]. Dense sampling is now routine, but it adds a dependency that hand-placed landmarks did not have: the quality of the correspondence, that is, how reliably the same dense point lands on the same anatomical location across specimens, now depends on the algorithm rather than on the analyst.

We used two tools to frame this study, ALPACA and DeCAL. They establish correspondence in different ways. ALPACA (Automated Landmarking through Point cloud Alignment and Correspondence Analysis; [9]) transfers the dense points of a reference specimen onto each target by point-cloud registration. It usually uses one reference, though several can be combined. It reduces the two surfaces to point clouds, aligns them rigidly, warps them onto each other with a deformable registration (a coherent-point-drift step), and records each reference point where it lands on the target. ALPACA is fast, runs on an ordinary computer, and needs no landmarks on the target specimens. But its correspondence is only as good as that registration, and a single reference can bias the result unless you combine several. DeCAL comes from the DeCA module (Dense Correspondence Analysis; [12]), which anchors the correspondence on the anatomical landmarks instead. It aligns the landmark configurations to a common mean by Procrustes superimposition, warps each specimen into the mean-shape frame with a thin-plate spline driven by those landmarks, and matches every atlas point to the nearest point on the specimen’s surface. It then carries the matches back to the specimen’s own frame with the inverse warp. We call this dense set DeCAL (DeCA Landmarking) to keep it separate from the module. The manual landmarks fix homology where we know it, and the dense points fill in between them by surface proximity. So the two methods need different things. ALPACA needs no landmarks on the targets and is quick to run. DeCAL needs anatomical landmarks on every specimen, but uses them to constrain the correspondence. Comparing them lets us ask two things: how many dense points to use, and how good the correspondence behind them needs to be.

These methods cost time and add complexity, so two questions matter to a working morphometrician.

- Q1. Do the dense semilandmarks recover shape that the standard anatomical landmarks miss, and how many points does it take? Denser sampling is only worth the effort if it adds information beyond the fixed landmarks, and if that gain does not saturate after the first couple of hundred points. Denser sampling is also not free statistically: thousands of coordinates, far more than the number of specimens, inflate the dimensionality of the data and can complicate the multivariate analyses.
- Q2. Does sliding help the biological question, or can it harm it? Sliding is near-universal practice [11], but its criteria, smoothness (bending energy) or among-specimen similarity (Procrustes distance), are geometric conveniences, not statements about homology, and the two criteria give different alignments [5]. If specimens really differ, a criterion that makes them look more alike could erase real variation as easily as it reveals correspondence.

The problem is that real specimens give no ground truth for correspondence. For a given semilandmark we do not know where its true homologue lies on the next skull, so we cannot grade a placement against truth directly. So we use a proxy: how well the workflow lets us reconstruct each specimen’s own scanned surface. As is standard in geometric morphometrics, we warp an initial reference template to the study’s mean shape, then mould this mean to each specimen using its own landmark positions from the GPA. We compare the reconstructed surface with the specimen’s real one. The better the landmarks capture the shape, the closer the reconstruction. This reconstruction-to-template error has been used before to weigh dense- sampling workflows against manual landmarks [13].

While the metric is useful, it is not equally good at detecting help and harm. The reconstruction score depends on the shape of the rebuilt surface, that is, how well it matches the real skull. It does not depend on which landmark sits at which point. But that is what correspondence is, and that is what sliding changes. Sliding sees only the landmark coordinates, not the mesh. It moves each semilandmark within a small estimated tangent plane. A good slide moves a point a short distance across the surface. This changes the correspondence but barely changes the shape, so the score hardly moves. The improvement is real but almost invisible. Sliding also does not know the surface, so the same move pushes the point slightly off it, and the score counts this as a small penalty instead of the correspondence gain it made. The score changes a lot only when sliding moves points far enough to distort the shape, as Procrustes-distance sliding does when it pulls every specimen toward a common mean. So the metric detects harmful sliding much better than helpful sliding. A result that shows no effect, or a small negative effect, on real data is therefore ambiguous: sliding may have done nothing, or it may have helped in a way the score cannot see.

We do this in two steps. First, we test Q1 and Q2 on 496 mouse skulls [14] with the surface-reconstruction metric, comparing workflows one specimen at a time. This gives a clear answer for Q1, which the metric measures well, but only a weak answer for Q2. Second, we settle Q2 with a simulation. Others have taken this same step of using controlled data when real specimens cannot give a ground truth. Some compare landmark-based and surface-based methods on real taxa, where the true differences stay unknown [15]. Others, closer to what we do, use simulated shapes whose differences are set in advance and score each method on how well it recovers them [16]. Our approach is complementary: we make synthetic skulls whose true point-to-point correspondence is known, so we can measure directly how far each semilandmark is from its true match. This measure does not depend on the surface at all. Together the two steps test sliding fairly, which neither could do alone.

## 2. Material and methods

### 2.1. Specimens

We used 496 laboratory-mouse skulls, the complete set imaged in the backcross of Maga et al. [14], in which an A/J × C57BL/6J F1 was backcrossed to A/J. That study’s quantitative-trait-locus analysis used only the 433 animals that also gave usable DNA markers; our morphometric analysis needs no genotypes, so we use all 496 imaged skulls. Mice were collected at postnatal day 28 and imaged by high-resolution micro-computed tomography (Skyscan 1076; 18 µm isotropic voxels, 55 kV, 0.5 mm aluminium filter). Cranial surfaces were reconstructed in 3D Slicer [17]. Following Maga et al. [7], we used an atlas-based segmentation to isolate the skulls as independent models. Each specimen has a high-density triangulated surface (of the order of one million triangles) and 55 fixed anatomical landmarks across the neurocranium, palate, and face (figure 1). No new animal procedures were undertaken for the present work (see Ethics).

**Figure 1.**
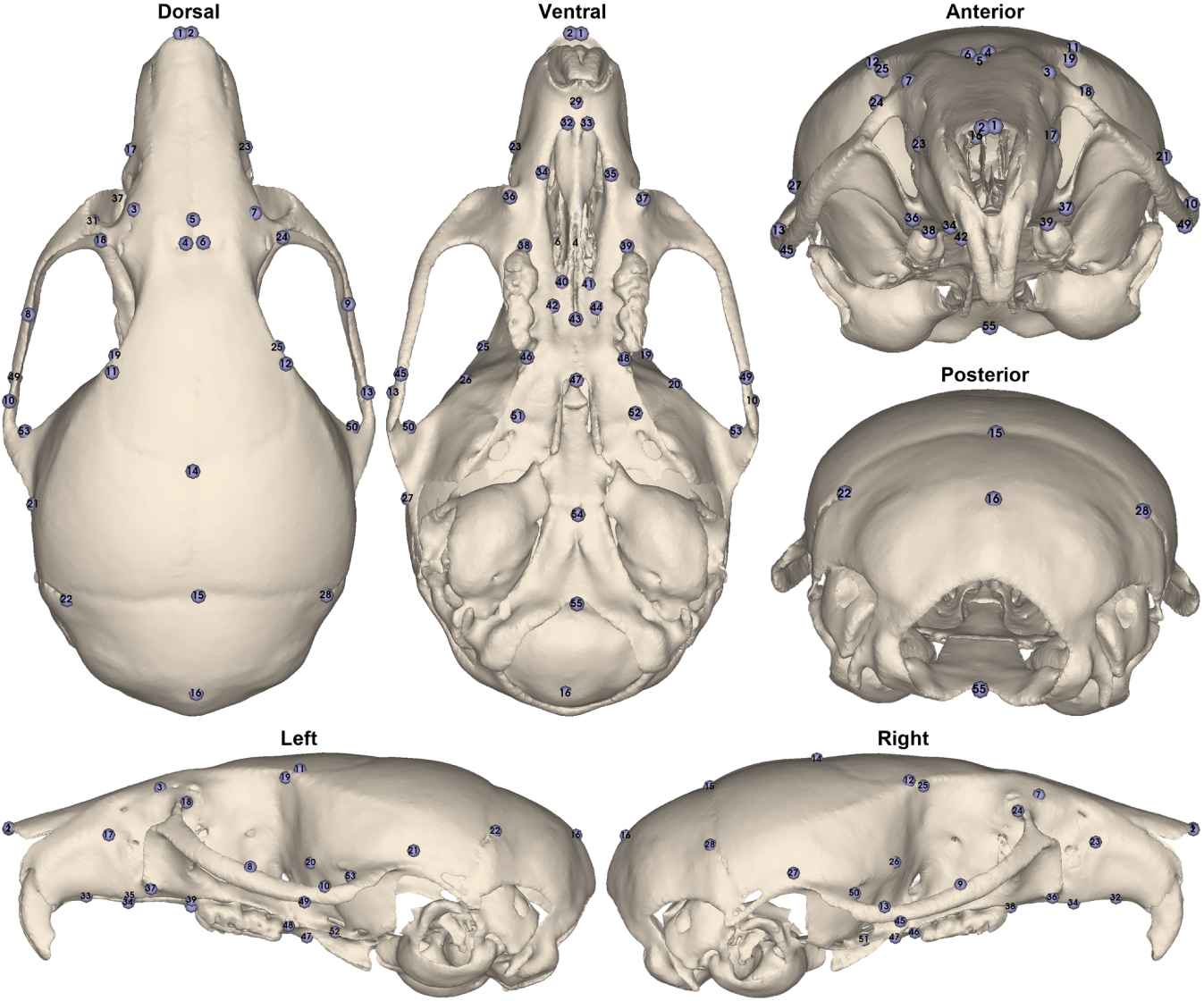
The 55 fixed anatomical landmarks used throughout the analysis of the real mouse skulls, shown on a representative specimen in dorsal, ventral, anterior, posterior, left lateral and right lateral views. The landmarks span the neurocranium, palate and face.

### 2.2. Reference sets: atlases and dense landmark workflows

We applied both workflows to the same 496 specimens (ALPACA and DeCAL; see the Introduction). For ALPACA, we generated an unbiased template from 30 randomly chosen skulls with ALPACA’s consensus template-generation routine [18]. We placed 1,002 points on that template with the SlicerMorph extension’s PseudoLMGenerator [17], and ALPACA’s single-template batch mode transferred them onto all 496 specimens [9]. For DeCAL, we gave the 496 specimens and their 55 fixed landmarks to the DeCA module, which built a mean atlas. From this atlas it reconstructed 1,002 dense semilandmarks and mapped them onto every specimen through the established correspondence [12]. So each workflow gives up to 1,002 corresponding dense points per specimen (figure 2), in that specimen’s own coordinate frame, together with a method-specific mean atlas surface. We use this atlas as the reconstruction template below.

**Figure 2.**
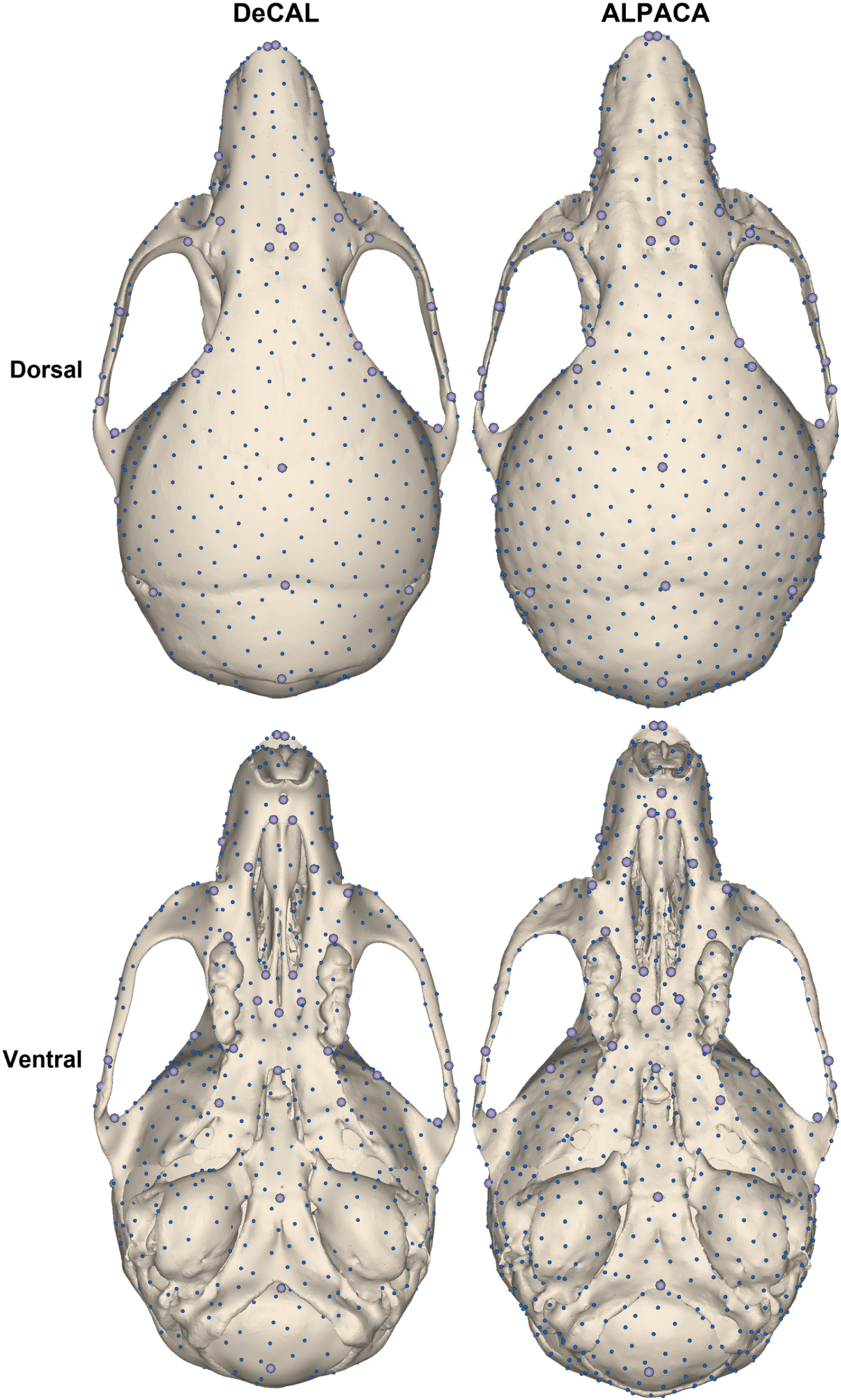
The dense point sets produced by the two workflows, each shown on its own mean atlas in dorsal (top) and ventral (bottom) views: the DeCAL semilandmarks on the DeCA mean atlas (left) and the ALPACA pseudo- landmarks on the ALPACA consensus template (right). Because each method builds its own atlas, the two point sets are not the same physical points.

### 2.3. Density ladder

To ask how many points are needed, we subsampled each method’s dense set to three nested densities, 250 ⊂ 500 ⊂ 1,000, by farthest-point sampling on the atlas. A specimen’s 250-point set is contained in its 500-point set, and its 500-point set in its 1,000-point set. We subsampled each method separately, because ALPACA’s points and DeCAL’s semilandmarks are not the same physical points.

### 2.4. Landmark workflows compared

We evaluated each density in two configurations: the dense points alone, and the dense points together with the 55 fixed anatomical landmarks. We compared both against a baseline of the 55 fixed landmarks only. This gives, per method, a fixed-only baseline, dense-only sets at 250/500/1,000, and dense-plus-fixed sets at 250/500/1,000. We scored every dense configuration both as placed and after sliding.

### 2.5. Sliding

We slid the semilandmarks with the new sliding routine in SlicerMorph’s GPA module, which follows the R/geomorph implementation as the de facto reference [19,20], minimizing either bending energy or Procrustes distance. In the dense-plus-fixed configurations we held the 55 fixed landmarks fixed and slid only the dense points; in the dense-only configurations all points slid. The sliding routine returns Procrustes-aligned, unit-scaled coordinates, so we returned each slid configuration to its specimen’s physical (millimetre) frame by a similarity fit on the fixed landmarks (or on all points, when there were no fixed landmarks) before reconstruction.

### 2.6. Surface-reconstruction metric

We scored each workflow by how well it reconstructs a specimen’s scanned surface. For a given specimen and workflow, we warped the method’s mean atlas surface by a thin-plate spline [21] so that the atlas landmarks met the specimen’s landmark positions for that workflow. The warped surface is the reconstruction. Reconstruction error is the signed closest distance from each reconstructed vertex to the specimen’s scanned surface (the nearest scan vertex; the roughly million-triangle scans make the vertices effectively continuous). We signed it by the scan’s surface normal, so positive values mean the reconstruction lies outside the true surface and negative values inside. We summarized each specimen and workflow by the root-mean-square distance (RMS, our primary error measure), and kept the mean absolute distance, the 95th percentile, and the signed mean (a net inflation or deflation bias) as secondary summaries. We computed the distance in one direction only, from the reconstruction to the scan. We considered a symmetric two-directional distance but did not use it, because the one-directional form is more conservative here: it is the less sensitive of the two to a landmark’s purely tangential shift along the surface, which is the very motion that sliding produces (Discussion).

The reconstruction template is each method’s mean atlas, which is itself a Procrustes consensus of the sample: a generalized Procrustes superimposition of the landmark configurations in DeCAL, and ALPACA’s iterative template generation. We reconstructed and scored every workflow, un-slid or slid, the same way: we warp this mean atlas by thin-plate spline onto the specimen’s landmark positions and measure the reconstructed surface against the specimen’s scan. The workflows differ only in which landmark positions we warp the atlas onto. An un-slid workflow uses the specimen’s landmarks as placed (in physical millimetres). A slid workflow first relaxes the specimen’s semilandmarks with the sliding generalized Procrustes analysis, then returns the relaxed positions to the specimen’s physical frame (above), and then does the same warp and measurement. So Procrustes superimposition underlies the reference mean in both cases; the sliding workflows just add the generalized-Procrustes sliding step that the un-slid workflows skip.

### 2.7. Comparisons and statistics

We compared workflows paired, specimen by specimen, across all 496 skulls. A single template cannot reproduce every idiosyncratic detail of a real skull, so all workflows share a common reconstruction floor, and what matters is the paired difference on top of that floor (lower RMS is better). We tested paired differences with the Wilcoxon signed-rank test and summarized them by the median paired change (with mean, standard deviation, and range) and by the proportion of the 496 specimens for which one workflow improved on another.

### 2.8. Software

We carried out all analyses in 3D Slicer [22] through its SlicerMorph [17] and DeCA [12] extensions: we did the point transfer (ALPACA) and the sliding generalized Procrustes analysis in SlicerMorph, and the dense correspondence (DeCAL) in DeCA. We generated the synthetic skulls with the SkullDeformExplorer module introduced here (see Data accessibility).

## 3. Results

All errors are the per-specimen surface-reconstruction RMS in millimetres, over the 496 skulls; lower is better. Each method warps its own atlas, so the fixed-landmark baseline differs between methods, and we make every comparison within a method and paired across specimens. Maps of the signed surface-reconstruction distance for 36 randomly selected specimens, for every method and condition, are provided in the electronic supplementary material (figure S1).

### 3.1. Reconstruction from the fixed landmarks alone

Warping to the 55 fixed anatomical landmarks reconstructed each skull’s surface to a mean RMS of 0.066 mm (SD 0.015; range 0.052–0.177) for the ALPACA atlas and 0.060 mm (SD 0.013; range 0.044–0.147) for the DeCAL atlas. Both baselines have a long upper tail, a few skulls that 55 landmarks reconstruct poorly (worst cases 0.15–0.18 mm). This is the variation the dense points have a chance to recover.

### 3.2. Q1: dense semilandmarks lower reconstruction error, and how many points are needed

Adding the dense semilandmarks lowered the reconstruction error sharply and for almost every specimen (table 1); we did no sliding in this experiment. The median improvement over the fixed baseline was 0.010–0.012 mm for both methods (Wilcoxon P < 10⁻⁸⁰ at every density), reached in 95.6% of specimens for ALPACA and 100% for DeCAL. The dense points also collapsed the between-specimen spread: the standard deviation fell from about 0.014 mm (fixed) to about 0.004 mm, and the poorly-reconstructed tail disappeared (maximum RMS from 0.18 to 0.067 mm for ALPACA). So they helped most the skulls the fixed landmarks served worst.

**Table 1.**
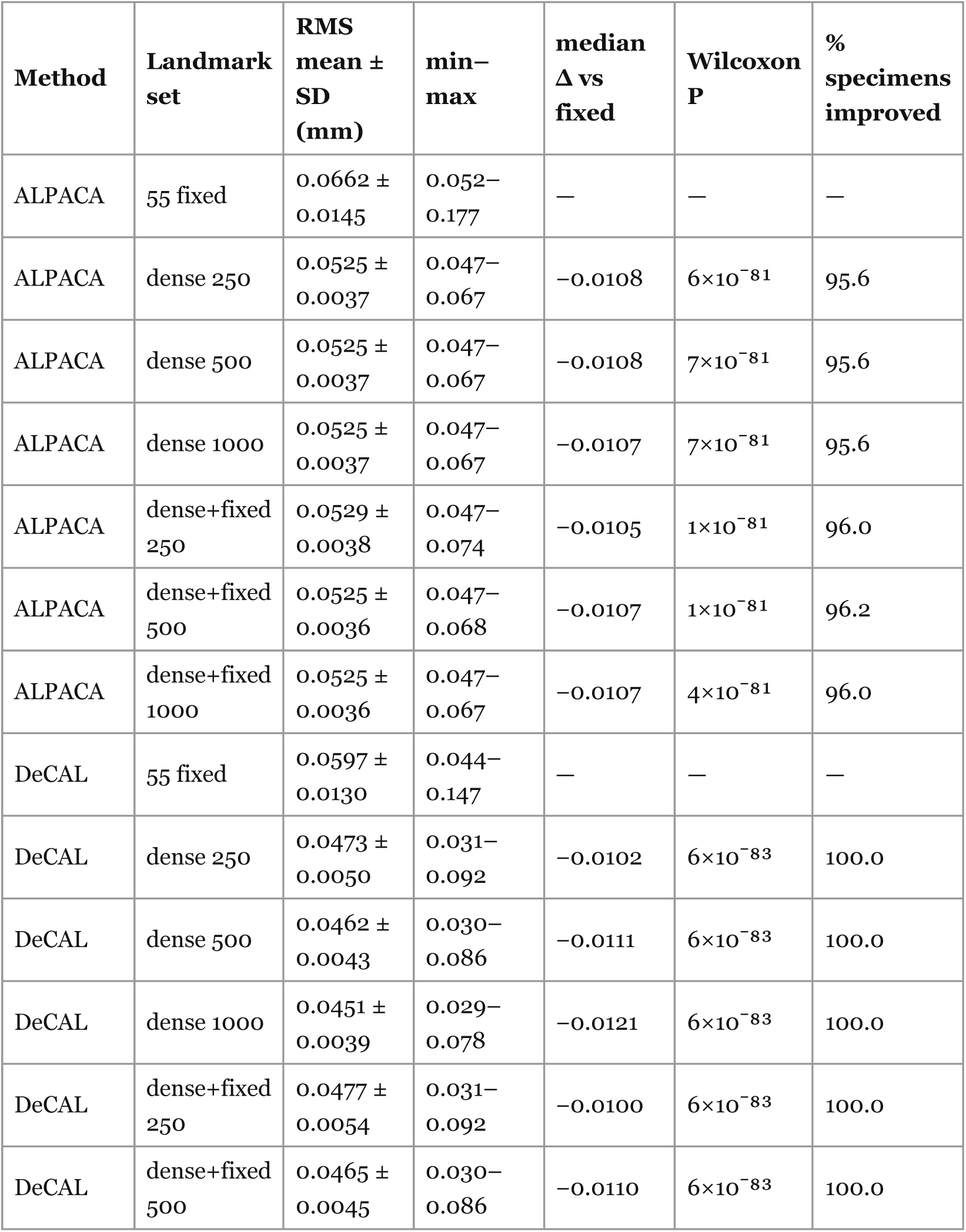
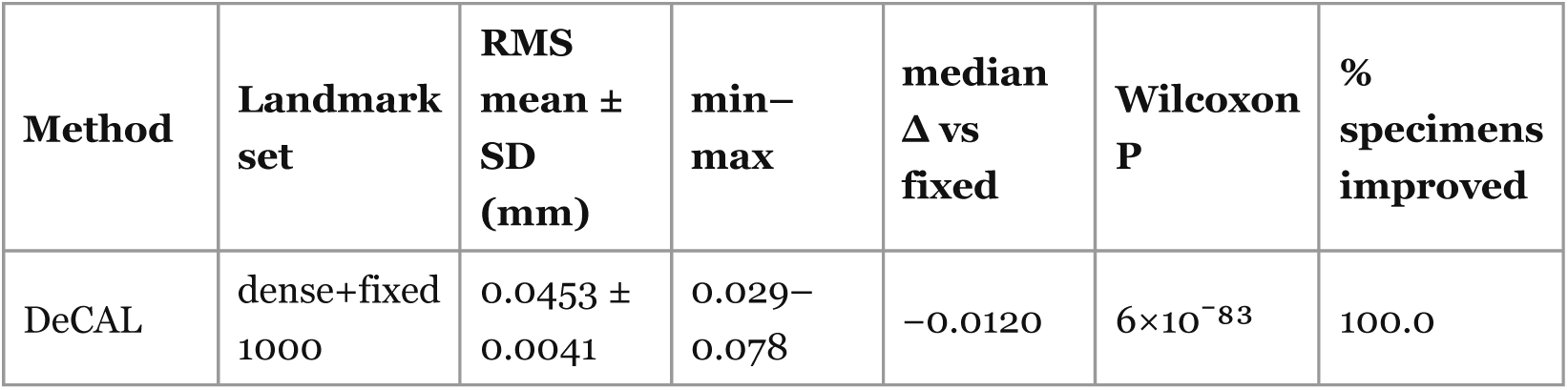
Reconstruction RMS by landmark set (no sliding); mean ± SD (min– max), and the paired change versus that method’s 55-fixed baseline.

| Method | Landmark set | RMS mean ± SD (mm) | min–max | median $\Delta$ vs fixed | Wilcoxon P | % specimens improved |
| --- | --- | --- | --- | --- | --- | --- |
| ALPACA | 55 fixed | 0.0662 ± 0.0145 | 0.052–0.177 | — | — | — |
| ALPACA | dense 250 | 0.0525 ± 0.0037 | 0.047–0.067 | –0.0108 | $6 \times 10^{-81}$ | 95.6 |
| ALPACA | dense 500 | 0.0525 ± 0.0037 | 0.047–0.067 | –0.0108 | $7 \times 10^{-81}$ | 95.6 |
| ALPACA | dense 1000 | 0.0525 ± 0.0037 | 0.047–0.067 | –0.0107 | $7 \times 10^{-81}$ | 95.6 |
| ALPACA | dense+fixed 250 | 0.0529 ± 0.0038 | 0.047–0.074 | –0.0105 | $1 \times 10^{-81}$ | 96.0 |
| ALPACA | dense+fixed 500 | 0.0525 ± 0.0036 | 0.047–0.068 | –0.0107 | $1 \times 10^{-81}$ | 96.2 |
| ALPACA | dense+fixed 1000 | 0.0525 ± 0.0036 | 0.047–0.067 | –0.0107 | $4 \times 10^{-81}$ | 96.0 |
| DeCAL | 55 fixed | 0.0597 ± 0.0130 | 0.044–0.147 | — | — | — |
| DeCAL | dense 250 | 0.0473 ± 0.0050 | 0.031–0.092 | –0.0102 | $6 \times 10^{-83}$ | 100.0 |
| DeCAL | dense 500 | 0.0462 ± 0.0043 | 0.030–0.086 | –0.0111 | $6 \times 10^{-83}$ | 100.0 |
| DeCAL | dense 1000 | 0.0451 ± 0.0039 | 0.029–0.078 | –0.0121 | $6 \times 10^{-83}$ | 100.0 |
| DeCAL | dense+fixed 250 | 0.0477 ± 0.0054 | 0.031–0.092 | –0.0100 | $6 \times 10^{-83}$ | 100.0 |
| DeCAL | dense+fixed 500 | 0.0465 ± 0.0045 | 0.030–0.086 | –0.0110 | $6 \times 10^{-83}$ | 100.0 |
| DeCAL | dense+fixed 1000 | 0.0453 ± 0.0041 | 0.029–0.078 | –0.0120 | $6 \times 10^{-83}$ | 100.0 |

How many points you need depends on the method (figure 3). For ALPACA the benefit had already plateaued at 250 points: the mean RMS was flat at 0.0525 mm across 250, 500 and 1,000 points, and the gain over baseline was the same (median −0.011 mm) at every density. For DeCAL the error kept falling as points were added, 0.0473, 0.0462 and 0.0451 mm at 250, 500 and 1,000, with the gain over baseline growing from 0.010 to 0.012 mm. DeCAL reconstructed better than ALPACA at every matched density (median difference 0.005–0.007 mm, P < 10⁻⁶⁵; ALPACA lower in only 3–9% of specimens), and that advantage widened with density (0.005 mm at 250 to 0.007 mm at 1,000). So DeCAL started lower and kept improving with denser sampling, while ALPACA saturated.

**Figure 3.**
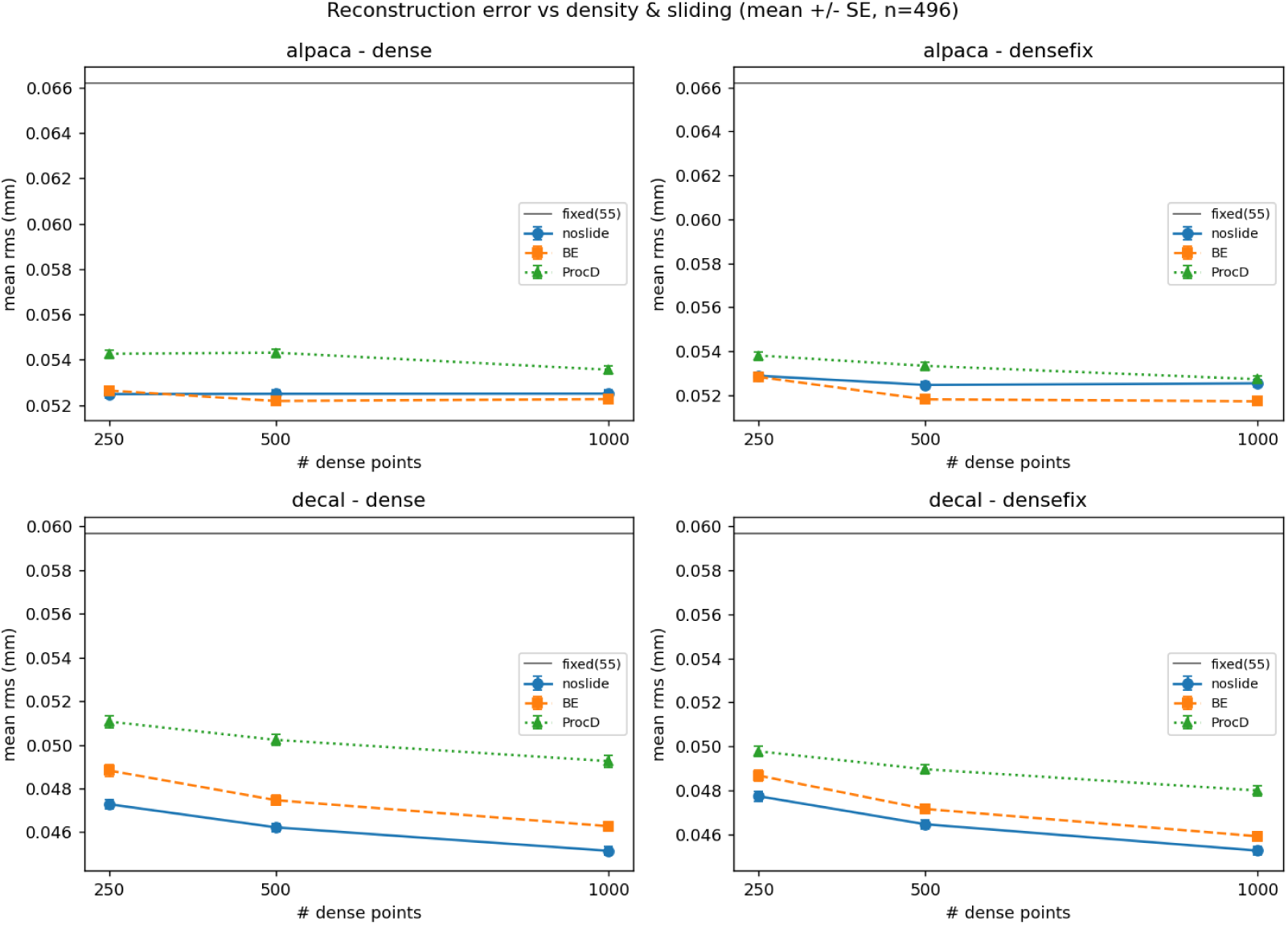
Mean reconstruction RMS (± standard error; n = 496) against dense point count (250/500/1,000), for each method (ALPACA, DeCAL) and landmark set (dense, dense-plus-fixed), shown for the un-slid configuration and after bending-energy and Procrustes-distance sliding, with the 55-fixed- landmark baseline for reference.

Adding the 55 fixed landmarks on top of the dense set made no practical difference: the paired change was at most 0.0005 mm and, for DeCAL, was slightly adverse (dense-plus-fixed improved on dense-only in only 3–9% of specimens). Once the dense points are present, the anatomical landmarks add nothing further to surface reconstruction as extra coordinates.

### 3.3. Q2: sliding does not improve reconstruction, and mostly harms it

Sliding the semilandmarks did not lower the reconstruction error in any consistent way, and its effects were an order of magnitude smaller than the dense-versus-fixed effect of Q1 (table 2, figure 4). Procrustes-distance sliding was harmful in every condition of both methods (median change +0.001 to +0.004 mm; it improved at most about 11% of specimens; P < 10⁻³⁰ throughout), which fits its pulling every specimen toward a common mean and erasing real differences.

**Figure 4.**
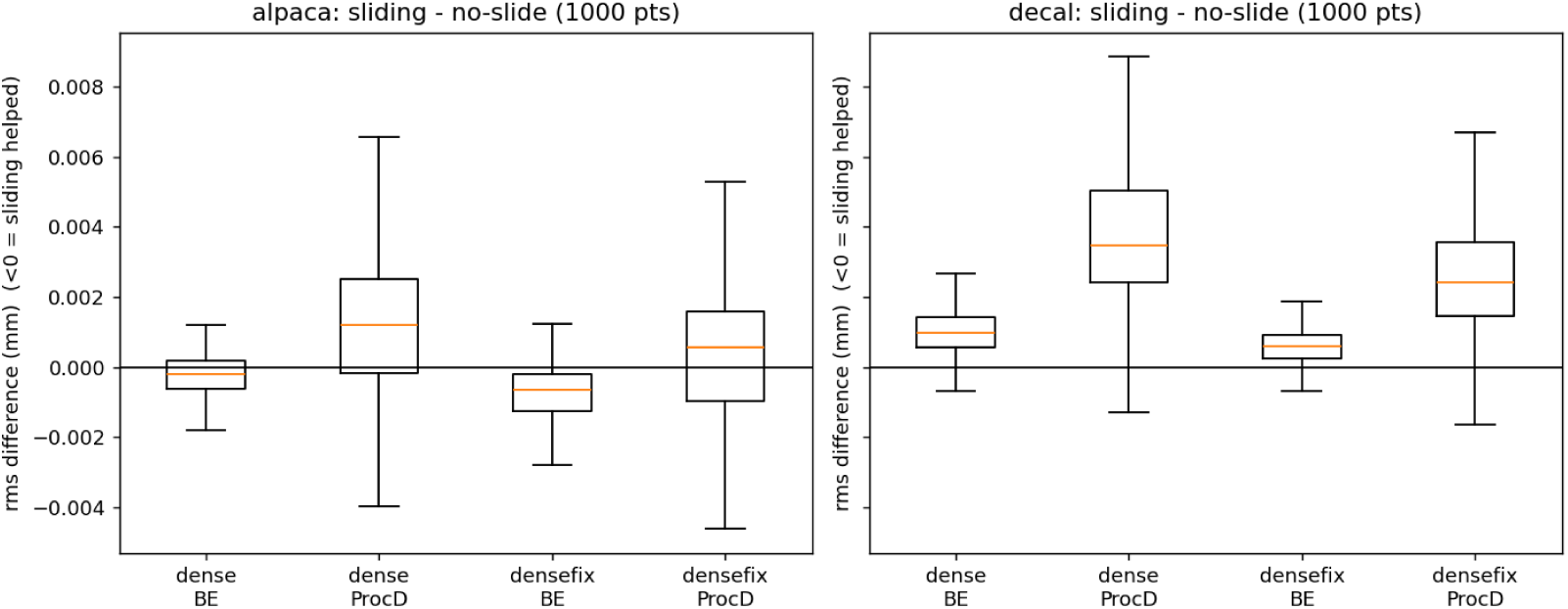
Paired within-specimen change in RMS from sliding at 1,000 points (slid minus un-slid; values below zero indicate sliding lowered the error), by criterion (bending-energy, Procrustes-distance) and landmark set (dense, dense-plus-fixed) for each method. Procrustes-distance sliding raises the error throughout, whereas bending-energy sliding lowers it only for ALPACA.

**Table 2.**
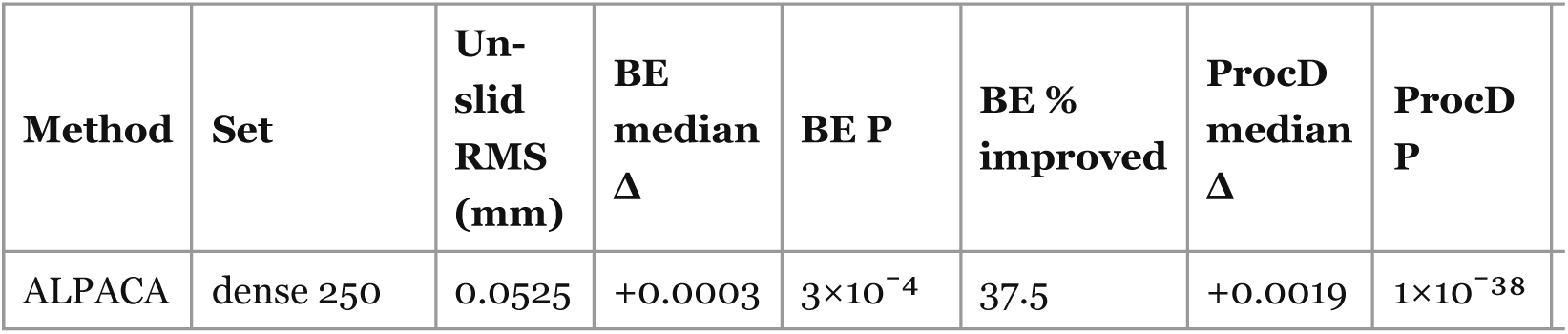
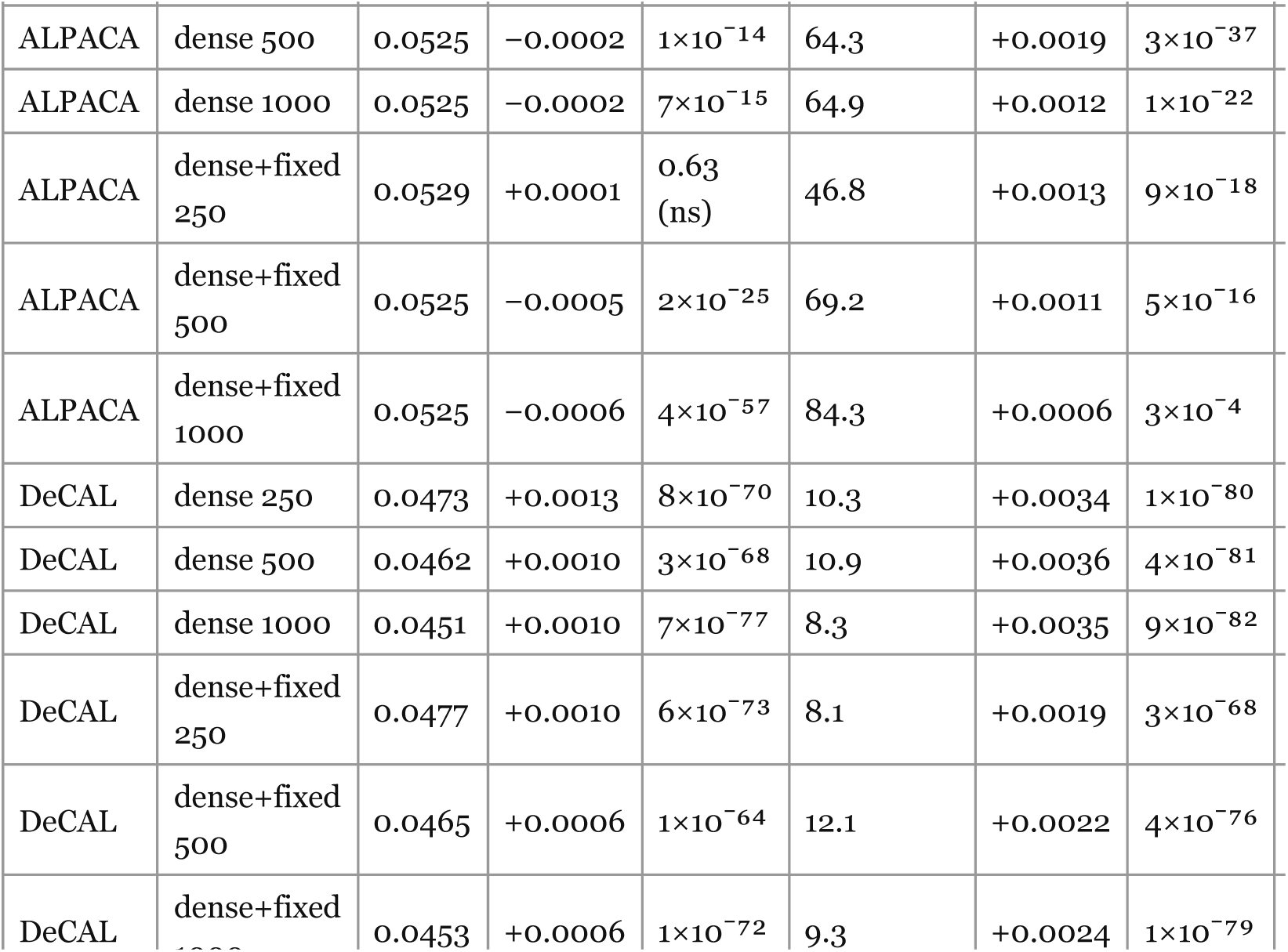
Effect of sliding, as the paired change (slid − un-slid) within each condition; a negative median favours sliding, and “% improved” is the fraction of specimens for which sliding lowered the RMS.

Bending-energy sliding depended on how good the correspondence already was. For DeCAL, whose correspondence is already high, it was harmful at every density (median +0.0006 to +0.0013 mm worse; improving only 8– 12% of specimens). For ALPACA, whose single-template correspondence is noisier, it gave a small improvement, clearest for the densest dense-plus- fixed set (dense+fixed 1000: median −0.0006 mm, 84.3% of specimens improved, P ≈ 4×10⁻⁵⁷) and more weakly for the higher-density dense-only sets (about 65% improved). Even where it helped, the effect was about 0.0006 mm, well below the between-specimen standard deviation and a tenth of the dense-versus-fixed gain. So on this surface metric the benefit of sliding, where it exists at all, is marginal.

So on real skulls the dense semilandmarks capture substantial shape that the fixed landmarks miss. This saturates by about 250 points for ALPACA but keeps improving to 1,000 for DeCAL. Sliding those points does not help in any way the surface metric can register, and Procrustes-distance sliding actively harms them. The one exception is a small bending-energy improvement for the noisier ALPACA correspondence, and it sits at the very edge of what this metric can resolve. Whether it is a real gain in correspondence is the question the synthetic experiment (Discussion) is built to answer.

## 4. Discussion

High-density semilandmark morphometrics, and the sliding at its centre, have been questioned. The concern is that Procrustes superimposition and bending-energy sliding can impose covariance the specimens did not carry, and that the risk is greatest in the dense configurations now most in use [23]. That debate is mostly about integration and modularity, that is, whether a module a study reports is real or an artefact of the fit. The concern is real, but integration and modularity are evolutionary-scale properties. Within a single population the modular structure is effectively fixed, so a study of one backcross such as ours does not depend on them. We also cannot settle that debate here, because there is no external standard to judge it against: whether one correspondence or one covariance structure is better than another cannot be read from the configurations themselves. It can only be judged against an outside criterion, such as agreement with an independent phylogeny. The present study is built on an explicit internal metric instead. Because the synthetic specimens have a known point-to-point homology by construction, we measure the distance from each placed or slid point to its true homologue. We then grade sliding by whether it moves each point closer to where it belongs, not by whether it gives a more plausible covariance structure. This lets us answer the sliding question directly, where the modularity debate can offer only indirect evidence.

The reconstruction metric has a limitation we should stress. It scores each landmark workflow by how well it rebuilds a specimen’s surface from a template, and that template is the sample’s own mean shape. So the measurement is not a fully independent, per-specimen quantity; it is anchored on a mean estimated from the same 496 skulls. Two things follow. In practice the anchor is stable here because the sample is large, so the mean is well estimated and the comparison of methods does not rest on a noisy template. More fundamentally, the group mean is the right level at which to judge a correspondence method: a method that really improves correspondence must improve the reconstruction of the whole sample relative to its mean, so an improvement measured this way is a population-level property, not a feature of any single skull. Even so, the results are statements about the methods, not about individual specimens.

This brings out the main difficulty with the second question. The metric answered Q1 without ambiguity, because adding dense points changes the shape of the reconstructed surface, which is what the metric was built to detect. Sliding is different. It moves a semilandmark tangentially, within a locally estimated tangent plane and with no knowledge of the mesh. It changes which point corresponds to which, not the shape of the reconstructed surface (Methods). So a surface metric is poorly suited to measuring what sliding does, and the Q2 effects show this: every sliding change was roughly 0.0006–0.004 mm, an order of magnitude below the dense effect and close to the resolution of the metric itself. Measuring the reconstruction-to-scan distance in one direction only (Methods) makes this worse by design: a one-directional distance is more forgiving of a purely tangential shift, so it understates any correspondence gain from sliding. We set the metric to under-credit rather than over-credit the very operation it is least able to see. The results are still informative. For ALPACA, whose single-template correspondence has the most placement error, bending-energy sliding lowered the reconstruction error in 84% of specimens at the highest density (P ≈ 4×10⁻⁵⁷), in the one case where there is correspondence error to correct; the same sliding harmed DeCAL, whose correspondence is already good. A benefit present in more than four-fifths of specimens is not noise; it is a real, population-wide effect. But its size on this metric is a fraction of a hundredth of a millimetre. This is a genuine gain the metric can detect in direction but cannot quantify: with 496 paired specimens even a small consistent shift is statistically clear, but the metric registers only the part of a tangential correction that changes the reconstructed shape, not the correction itself. So the surface metric can tell us that sliding helps when correspondence is poor and harms it when correspondence is already good, but not by how much, and it would miss any subtler benefit.

To answer the question properly we have to measure correspondence itself, not the surface, which means knowing where each semilandmark’s true homologue lies on every specimen. Real scans cannot give us this, but a controlled simulation can. We built a new 3D Slicer module, SkullDeformExplorer (archived at Zenodo; see Data accessibility). It builds a population of synthetic skulls by deforming a single template mesh with an a priori statistical shape model. Each specimen is the template warped by a random draw over the principal modes of real mouse-skull shape variation, plus an anisotropic change in size. We derived the principal modes from an independent reference sample of 62 laboratory-mouse skulls (the *Mouse_Models* data repository; [7]). We generated an unbiased consensus surface from these skulls with ALPACA’s iterative template-generation routine [18], placed 1,160 pseudo-landmarks on the consensus with SlicerMorph’s PseudoLMGenerator [17], and transferred them onto all 62 skulls with ALPACA [9]. A generalized Procrustes analysis of the 1,160-point configurations, followed by principal component analysis, gave the mean shape and the principal components from which each synthetic specimen is drawn; we used the ten leading components for generation. Because every specimen is a deformation of the same mesh, its vertices stay in exact correspondence across the whole population, so we know the true homologue of every point by construction. Onto these known-correspondence skulls we add realistic placement error by placing the dense semilandmarks with the same routes we used on the real data, not by any idealised rule. So the points carry exactly the correspondence error those methods make in practice: they lie on the surface but sit tangentially off their true homologue, which is exactly the error sliding is meant to repair. The evaluation then measures what the real-data metric cannot: the distance from each placed semilandmark to its true homologue, before and after sliding. Because we read this distance from the known vertex correspondence and not from the surface, a slide that moves a point toward its homologue is credited in full, and a slide that moves it away is penalised. So the tangential motion the surface metric cannot see becomes the quantity we measure directly.

For this trial the module made 500 skulls from a single 84,216-vertex mouse-skull template. Each is an independent draw over the ten leading modes of mouse-skull shape (uniform within ±3 standard deviations), with overall size drawn log-normally over a comparable range and an anisotropic rescaling of the axes. We left the surfaces smooth so that every placement error came from the correspondence methods and not from surface texture. The resulting morphospace is broad. Measured on the anatomical landmarks, the mean Procrustes distance to the consensus was 0.081 (SD 0.028; range 0.033–0.189), and centroid size ranged from 33.6 to 57.8 mm (mean 44.3, SD 4.9; coefficient of variation 11.1%). For comparison, the 55 fixed landmarks of the real backcross give a mean Procrustes distance to the mean of 0.027 (SD 0.004) and a centroid-size coefficient of variation of 2.7%. So the synthetic skulls cover, and go beyond, the shape and size variation of the real specimens, and the sliding test is not confined to a narrow range of form. The module can also add fine texture detail, but we did not use this. We placed the dense semilandmarks on all 500 skulls by both routes we used on the real data: ALPACA, transferring roughly a thousand points of the undeformed template onto each specimen by point-cloud registration without landmarks, and DeCAL, deriving a comparable set through each specimen’s fixed landmarks. This lets us judge the noisier and the anchored correspondence apart. We slid each placed set using 250, 500 and 1,000 points by both criteria, each method in the condition in which it is actually used. For ALPACA, which places no landmarks, all dense points slid freely; this is the case the real data left open, where correspondence is noisiest and sliding has the most room to help. For DeCAL, whose correspondence is built from the fixed landmarks, we held those fixed landmarks as anchors and slid only the dense semilandmarks, exactly as in the dense-plus-fixed analysis of the real skulls. We then recorded, for every semilandmark on every skull, the distance to its true homologue before and after sliding, summarised each specimen by its mean, and compared the paired values across the 500 with the Wilcoxon signed-rank test. This scores the tangential correction against the known vertex rather than the reconstructed surface, so it credits or penalises the move in full.

We show the sliding comparison for ALPACA. Its landmark-free correspondence is the noisy case where sliding has room to act. DeCAL’s landmark-anchored correspondence is already good, so sliding it here only repeats the do-not-slide result we saw on the real skulls (table 2). So ALPACA is the informative test of whether sliding recovers true homology. On these synthetic skulls the placed points already lay close to their true homologues before any sliding: for ALPACA, a mean of about 0.20 mm from the truth (table 3), which shows how accurately point-cloud registration transfers a template onto smooth surfaces. Bending-energy sliding then moved ALPACA’s points measurably closer to their homologues, and more so as they grew denser. At 250 points the change was neutral to slightly adverse (median +0.003 mm, 43% of specimens improved). At 500 and 1,000 points it became a consistent gain (median −0.007 and −0.009 mm; 68% and 82% of specimens improved; P < 10⁻²⁰). This is the result the surface metric could not resolve. Read against known homology rather than the reconstructed surface, bending-energy sliding of a noisy, landmark-free correspondence really does recover correspondence: the benefit is population-wide, it grows with density, and the 82% of specimens improved at 1,000 points closely matches the 84% the surface metric registered, but only faintly, on the same effect in the real skulls. Procrustes-distance sliding did the reverse at every density, carrying ALPACA’s points about 0.20 mm further from their homologues and improving not one specimen (P < 10⁻⁸⁰). This is what a criterion does when it pulls configurations toward a common mean: it overwrites the differences under study.

**Table 3.**
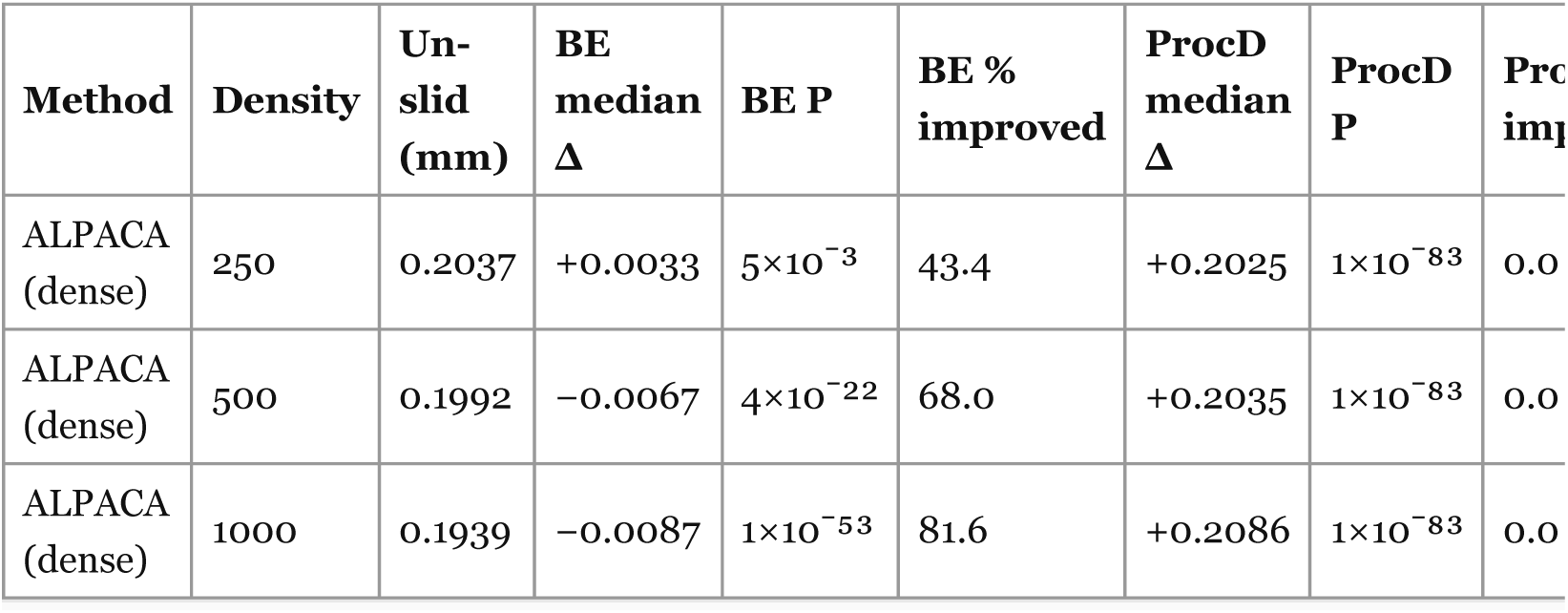
Sliding on the synthetic skulls, scored as the distance from each dense point to its true homologue (mm): the un-slid distance and the paired change (slid − un-slid) within each condition. A negative median favours sliding, and “% improved” is the fraction of the 500 specimens for which sliding lowered the distance to homologue. Only ALPACA is shown; it is slid dense-only, as it places no landmarks.

These known-correspondence skulls also settle a question the real data raised but could not answer: how much anatomical-landmark coverage is enough that sliding the dense points no longer helps? This is a question for DeCAL, not ALPACA, because DeCAL builds its dense correspondence from the anatomical anchors. So thinning those anchors shows where the correspondence passes from loose enough to benefit from sliding to good enough to be harmed by it. ALPACA has no anchors, so it cannot show this. We held DeCAL’s dense sampling fixed at about 500 points and thinned the anatomical anchors from 34 landmarks down through 27, 20, 13 and 8. We chose each subset by farthest-point sampling so that, however few, it still spanned the whole skull, and we slid the dense points with that subset held fixed. The question began as whether one particular 27-landmark set was still enough, so we ran the whole ladder rather than that single count, to locate the threshold itself. To separate the number of anchors from their spatial coverage, we added two count-matched controls in which the same 27 or 13 landmarks were clustered on a single region of the skull. The anchor positions are error-free throughout; we varied only their number and placement, not their accuracy.

The benefit of bending-energy sliding fell steadily as we added anchors, crossing from help to harm between about 20 and 27 well-spread landmarks (table 4). With 8 anchors it drew the dense points closer to their true homologues in 66% of specimens (median −0.0033 mm); with 13, in 60%; at 20 and 27 the effect was indistinguishable from zero (51% and 46% of specimens; P = 0.72 and 0.12); and by 34 it was a reliable harm (39%; P = 9×10⁻⁸). So the coverage-preserving 27-landmark set that prompted the question already sits in the do-not-slide regime. The clustered controls show why: 27 landmarks bunched on one region left the correspondence much poorer (un-slid distance 0.196 mm, against 0.153 mm for the well-spread 27), and sliding then clearly helped it, in 69% of specimens. So a well-placed 27 and a badly-placed 27 fall on opposite sides of the threshold. It is the spatial coverage of the anchors, not their raw count, that sets whether the correspondence is already good enough that sliding can only disturb it. Procrustes-distance sliding harmed the configuration at every rung of the ladder, improving no specimen, as it did everywhere else.

**Table 4.**
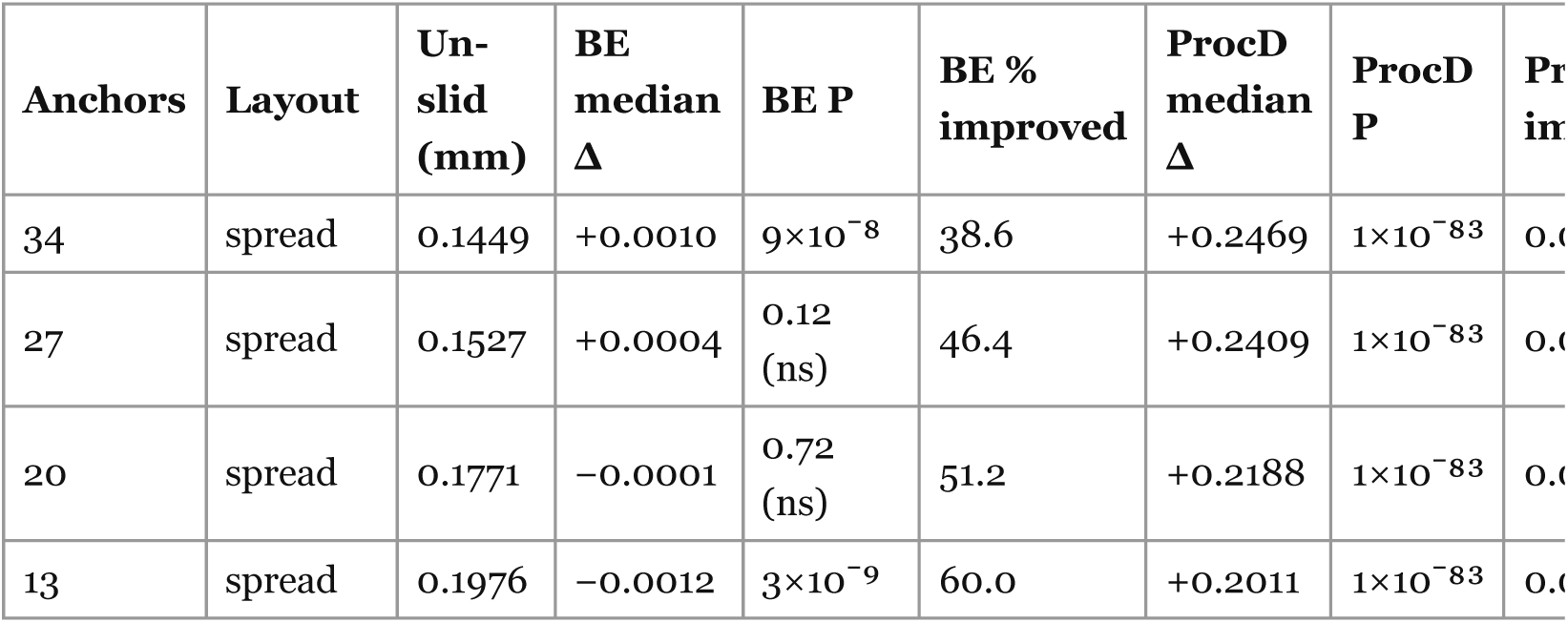
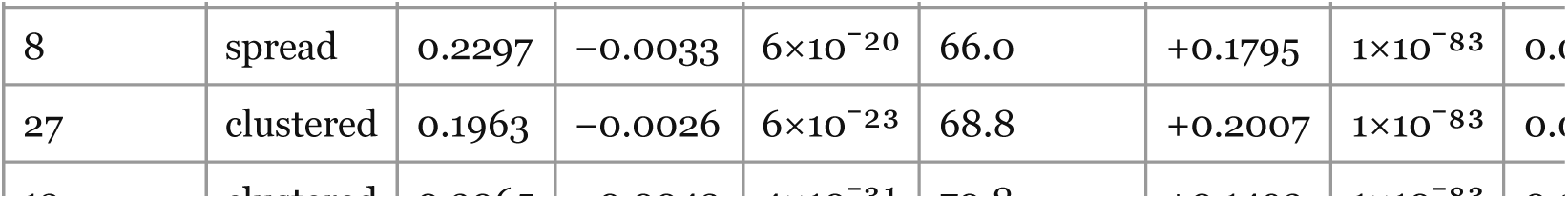
Fixed-landmark ablation on the synthetic skulls: the distance-to- homologue effect of sliding as the number and coverage of (error-free) anatomical anchors is thinned, with the dense sampling held at ∼500 points. The “spread” subsets span the whole skull by farthest-point sampling, and the “clustered” subsets of the same count are confined to one region. Un-slid distance and the paired change (slid − un-slid); a negative median favours sliding.

So what should a practitioner do? For the single-population studies where automated dense landmarking is normally used, our results give a few simple rules. Each one depends on how good the correspondence already is.

If the dense points come from DeCAL, the decision depends on the anatomical landmarks used to build it. If those landmarks are many and well spread over the skull (the 55 used in the real data, or about 20 or more well-spread points), do not slide. The landmarks have already set the correspondence, and bending-energy sliding only makes it worse: it raised the reconstruction error in every condition of the real data (table 2) and never lowered it. If the landmarks are few, or clustered in one area (below about 20 well-spread points), the correspondence is weaker, and bending-energy sliding then helps, moving the dense points closer to their true matches (table 4).

If the points come from ALPACA and the study has no anatomical landmarks at all, just a dense set of pseudo-landmarks, then bending-energy sliding is the right choice. This is the one case where the correspondence is loose enough that sliding improves it, moving points closer to their true matches in most specimens, and more so as the points get denser.

If you have anatomical landmarks and only want to add points between them, do not slide an ALPACA set. Use DeCAL from the start. It turns those landmarks into a denser, higher-quality sampling that keeps improving as you add points, whereas ALPACA’s single-template correspondence had already stopped improving.

Finally, never use the Procrustes-distance criterion to slide. In every condition of both methods, on real and synthetic skulls alike, it pulled specimens toward a common mean and moved points away from their true positions, erasing the very variation the analysis is meant to measure. A last point concerns the anatomical landmarks themselves. Placing well-defined landmarks by hand is tedious, but our results show it is not wasted effort and not an obstacle to dense analysis. Those landmarks are what anchor a dense correspondence: they let DeCAL place points more accurately than the landmark-free route, and a well-spread set of them removes the need to slide at all. Good anatomical landmarks do not hold dense morphometrics back; they are what make it reliable.

These rules come with a caution. They are based on one population: a single backcross of laboratory mice, plus synthetic skulls covering its range of variation. But this is the setting where automated dense landmarking is used most often, a template transferred between specimens that are already broadly similar, so it is the common case, not a special one. When a sample spans larger differences, such as across genera or a wide phylogenetic range, correspondence is harder to establish. There, both the value of dense sampling and the balance between the benefit and harm of sliding may change, and the rules above should be tested rather than assumed. For the usual single-population study, the advice is simple: match the sliding decision to the correspondence you have. Relax a poor one, leave a good one alone, and never slide toward the mean.

These limits also point to future work. The approach used here, building specimens whose true correspondence is known and then testing a workflow against that truth instead of a proxy, is not limited to single populations. The same idea could be applied at the macroevolutionary scale, where integration and modularity actually appear, and could give those much-debated analyses an internal check they currently lack, since today they can only be judged against an external proxy such as agreement with a phylogeny. Such an extension does not need a new statistic. It needs the raw material: the full scanned surfaces together with their landmarks, openly shared. This is what is most often missing. Many high-density studies release only the processed output, Procrustes-aligned coordinates or the covariance summaries computed from them. These are enough to reproduce a published figure, but not enough for the kind of ground-truth re-analysis done here, which has to start from the surfaces themselves. The field would be better served by archiving the full datasets, meshes and landmarks alike, in open, non- proprietary formats, than by the derivatives, or the closed formats, that usually stand in their place. Only open primary data let a method be tested again, rather than just re-displayed.

## Ethics

The micro-computed-tomography images analysed here were generated in a previous study [14]; no new animal procedures were undertaken for the present work. All animal protocols in that study were approved by the Institutional Animal Care and Use Committee of the University of Washington (protocol #2688-07).

## Data accessibility

The 496 mouse-skull surface models and their anatomical landmarks are available at the Open Science Framework (https://osf.io/sn64d/), in the “sliding” folder. The SkullDeformExplorer module used to generate the synthetic skulls, part of the MorphoSim extension, is archived at Zenodo (https://doi.org/10.5281/zenodo.21515663). The dense landmarking, sliding and correspondence analyses of the real skulls were run with the openly available, published 3D Slicer extensions SlicerMorph [17] and DeCA [12]. The reference sample used to build the shape model is the *Mouse_Models* data repository [7]. Supporting information, including figure S1, is provided as electronic supplementary material.

## Declaration of use of AI

During the preparation of this manuscript the author used a large language model (Claude, Opus 4.8; Anthropic) to assist in developing and testing the analysis and figure-generation scripts and in drafting and editing text. The author reviewed, tested and validated all AI-assisted code against known cases, checked all text, and takes full responsibility for the content of this manuscript.

## Authors’ contributions

A.M.M.: conceptualization, data curation, formal analysis, funding acquisition, investigation, methodology, software, validation, visualization, writing—original draft, writing—review and editing. The author read and approved the manuscript and agrees to be accountable for all aspects of the work.

## Acknowledgements

The author thanks the SlicerMorph and 3D Slicer communities for the open-source tools that made this work possible.

